# Monogamy in a moment: how do brief social interactions change over time in pair-bonded zebra finches (*Taeniopygia guttata*)?

**DOI:** 10.1101/2020.06.18.160051

**Authors:** Nora H. Prior, Edward Smith, Robert J. Dooling, Gregory F. Ball

## Abstract

Research on monogamy has largely focused on marked behaviors that are unique to pair bonded partners. However, these marked behaviors represent only a subset of the pair-directed behaviors that partners engage in; the influence of pair bonding on mundane or subtle social interactions among partners remains largely unknown. In the current study, we describe the changes that occur during brief social reunions (or greets) over the course of pair bonding in zebra finches. We quantified pair-directed behavior during five-minute reunions from three stages of pair bonding: initial pairing (between 4-72 hrs), early pairing (1-2 weeks) and late pairing (>1 month). These social interactions were operationalized in multiples ways. First, we quantified the overall activity levels (call and movement rates) for both the male and female. Overall, females were more active than males, but for both males and females calling activity was highest during the initial timepoint (between 4-72 hrs post-pairing). We quantified behavioral coordination between partners in two ways, 1) similarity in call and movement rates between partners, and 2) temporal synchrony between calls and movements (via sliding correlation coefficients of time-stamped calls and movements). Overall there were no effects of pairing on behavioral coordination. Finally, we used principal component analyses to disentangle behavioral coordination from the activity levels of the male and female. These results contribute to a growing line of evidence that male and female zebra finches differentially contribute to social dynamics and highlight the influence of pair bonding on the development of social dynamics. Behavioral coordination is clearly important for marked interactions (e.g. duetting, courtship displays and biparental care). Our results raise the question of what the roles of such mundane social interactions are in monogamous partnerships.

## Introduction

In monogamous species, the formation and maintenance of a pair bond is necessary for the successful rearing of offspring (Lack, 1968; Reichard & Boesch, 2003). The majority of research on the mechanisms underlying pair bonding has focused on marked, pair-specific behaviors and interactions of partners (Aragona et al., 2006; Soma & Garamszegi, 2015; Wachtmeister, 2001; Young et al., 2011; Young & Wang, 2004). This approach has revealed remarkable conservation in the mechanisms of pair bonding across taxa (Bales et al., 2007; Donaldson & Young, 2016; O’Connell & Hofmann, 2012; Walum & Young, 2018).

Highly-marked behaviors and interactions are particularly evident during courtship and initial pair-bond formation. For example, in prairie voles (*Microtus ochrogaster*), the establishment of a pair bond is clearly marked by the development of selective preference/affiliation for a mate (Resendez et al., 2016; Williams et al., 1992; Young et al., 2011). This marked selective attachment has been used to identify the mechanisms underlying pair formation. However, these marked behaviors represent only a subset of those that partners engage in, and more detailed understandings of partner interactions are needed in order to further elucidate mechanisms of pair bonding. During partner preferences tests, the pattern of approach and proximity time between partners continues even after initial establishment of the pair bond, with the focal individual spending more time with their partners at later stages of pair bonding (Scribner et al., 2020). Furthermore, important long-term impacts of pair bonding on the brain and behavior of voles are revealed by differences in approach of an individual to its mate versus a stranger (Scribner et al., 2020). Even in the most commonly studied model systems, remarkably little is known about the consequences of pair bonding on mundane or subtle features of partner interactions.

Whereas monogamy is rare in mammals (<4% of species), the vast majority of birds form some type of monogamous partnership (∼90% of species) (Reichard & Boesch, 2003). Across bird species there is considerable variation in the phenotype of monogamous bonds. Monogamous bonds vary in how long they are maintained; they may be transient, lasting only a season, or life-long (Black & Hulme, 1996). As with prairie voles, the majority of studies focus on marked features of pair bonding (e.g. partner preference, proximity time, allopreening and clumping) (Kenny et al., 2017; Prior et al., 2013; Smiley et al., 2012; Tomaszycki & Adkins-Regan, 2005). However, the importance of brief social interactions has also been described in numerous species: at the nest, brief social interactions appear to be essential for the active coordination of parental duties between partners (I. C. A. Boucaud et al., 2016; Mariette & Griffith, 2012, 2015; van Rooij & Griffith, 2013). This raises the question more broadly of how subtle features of social interactions are shaped by pair bonding.

Here we describe the effect of pair bonding on brief social interactions in monogamous zebra finch pairs. Zebra finches maintain sexually monogamous life-long pair bonds, are non-territorial, and are highly gregarious. Interestingly, traditional partner preference paradigms may fail to demonstrate selective preference for the partner (Prior et al., 2013), although other behavioral assays clearly show that many aspects of pair-directed behavior are reserved for or more common between partners than familiar conspecifics (Fernandez et al., 2017; Gill et al., 2015). Additionally, intra-pair calling dynamics, across multiple contexts, appear to be important behavioral components of the zebra finch pair bond (I. Boucaud et al., 2016; Pietro Bruno D’Amelio et al., 2017a; Elie et al., 2010; Fernandez et al., 2017; Gill et al., 2015).

In the current study, we operationalize brief social reunions (both calls and movements) of pairs over the course of pair bonding: initial pairing (between 4-72 hrs), early pairing (1-2 weeks) and late pairing (>1 month). First, we quantified the overall activity levels (call and movement rates) for both males and females. Second, we used two approaches to estimate the coordination of the activity between partners, including quantifying the similarity in call and movement rates between partners as well as quantifying the sliding correlation coefficients for time-stamped calls and movements (a measure of temporal synchrony). Finally, we used principal component analyses to disentangle behavioral coordination from the activity levels of the male and female.

## Materials and Methods

### Subjects and establishment of pairs

Twenty adult zebra finches (5-6 months old) were used in this study (10 females and 10 males). Throughout the study, zebra finches were housed with *ad libitum* seed, water and grit on a 12L:12D light cycle. This same cohort of zebra finches was also used for a subsequent experiment (Prior et al., 2019).

Prior to pairing, zebra finches were housed in same-sex flocks. In order to provide the opportunity for pairs to freely form, birds were moved to mixed-sex flocks for 72 hours. Providing individuals with the opportunity to freely form monogamous bonds is important as forced pairing can be associated with lower pair fecundity (Griffith et al., 2017). Thus, birds were housed in mixed-sex flocks with either a male- or female-biased sex ratio (2 females with 3 males or 3 females with 2 males). Pair bonding was assessed visually each day; occurrences of selective affiliative behaviors (i.e. clumping, allopreening, and coordinated preening) between individuals were scored during 5 min behavioral observations. After 72 hours, birds were removed from mixed-sex flocks and housed with their mates for the duration of the study. Note that it is typical for not all birds in mixed-sex flocks to form pair bonds (Scalera & Tomaszycki, 2018; Smiley et al., 2012). Here we identified four clearly-established pairs (paired). Another four pairs were selected from the same mixed-sex flocks of birds that showed little evidence of pairing. The last two pairs were composed of birds unfamiliar with each other. Thus, we established 10 zebra finch pairs with varying extents and patterns of prior experience. We predicted that prior experience would affect behavioral coordination in our reunion paradigm. Thus, despite that there were few birds in each group, we used prior experience as a factor in our later analyses (Paired N=4, Weakly Paired N=4, Force Paired N=2). All pairs, regardless of prior experience, were seen being highly affiliative in the home cage (and were not seen interacting aggressively). Additionally, after this experiment, all pairs attempted to breed (nest building and/or egg laying) and 8 out of 10 pairs successfully fledged chicks.

### Experimental Design

A timeline of the behavioral recordings is presented in Figure 1. We recorded brief social interactions from each pair nine times over the course of the first month of pairing. These nine recordings were not evenly distributed over the course of the month, but were instead situated within periods of the pair bonding process commonly described in the research. Pairing can be conceptually divided up into three stages: a brief courtship phase, a short pair formation phase, and an indefinite pair maintenance phase. Although these stages are commonly referenced (Prior & Soma, 2015; Resendez et al., 2016; Scribner et al., 2020; Smiley et al., 2012), what distinguishes these stages and how long they last is unclear. In general, pair maintenance encompasses anything that occurs after the establishment of a pair bond. In zebra finches, pair bond formation can take up to two weeks; however, it is typically assumed to occur much more quickly (on the order of hours to days) (Zann, 1996).

**Figure 1:**
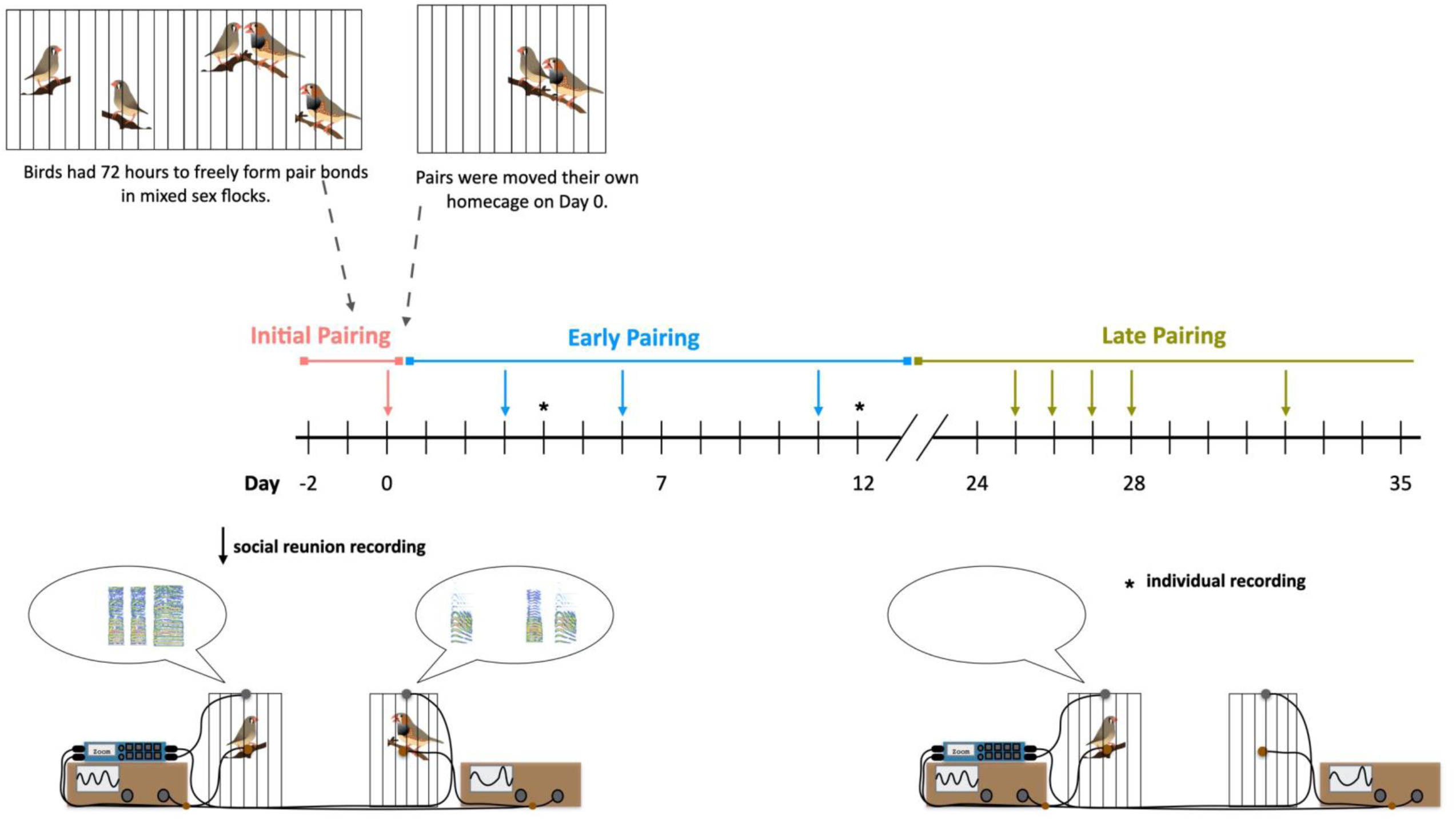
Diagram of the timeline of initial pair formation and behavioral recordings during the first month of pairing. All birds were given the opportunity to freely form a pair bond in mixed sex flocks over the course of 72 hours. The first social reunion was recorded on the day that pairs were moved from mixed-sex flocks to a new home cage with their partner (“*initial pairing*”). At this timepoint, pair had been removed from flock for 4-72 hours. At the time of pairing, four pairs from the mixed-sex flocks clearly engaged in selective pairing behavior, four pairs were formed from birds that had been together in mixed-sex flocks but were not obviously pair bonded, and two pairs were composed of individuals that had not been in the same mixed-sex flocks and had no prior experience. We recorded reunion behavior from 3 timepoints during the following two weeks (“*early pairing*”), and from 5 timepoints during a late stage of pairing (>1 month post pairing) (“*late pairing”)*. Additionally, as a control, two recordings were made of each individual alone in the testing room (individual recordings). A schematic illustration of social reunion behavior paradigm is shown in the bottom right; redrawn from Prior et al., 2019). Briefly, behavior was recorded using four channels of a Zoom recorder (F8): movement was recorded from a piezo sensor attached to the perch of a smaller cage (indicated via oscilloscope), and acoustic behavior was recorded using tie clip microphones (indicated via spectrogram).

Here we recorded the first social reunion on the day that pairs were moved from mixed-sex flocks to a new home cage with their partner (“*initial pairing*”). At this timepoint, all pairs had been given the opportunity to engage in courtship behaviors and copulate; however, depending on how they were paired (*described above*) this time ranged from 4-72 hours together. Thus, we would predict that a mix of courtship and pair formation could be occurring at this timepoint. Next, we recorded reunion behavior from 3 timepoints during the following two weeks (“*early pairing*”). By two weeks, we would expect all pair bonds to have become established. The majority of research on pairing does not extend past 2-3 weeks post-pairing (Scalera & Tomaszycki, 2018; Smiley et al., 2012; Tomaszycki & Adkins-Regan, 2005; Tomaszycki & Zatirka, 2014), and anything beyond that is typically considered pair maintenance (Prior et al., 2013; Scribner et al., 2020; Tomaszycki & Adkins-Regan, 2006). Finally, we recorded reunion behavior at 5 timepoints during a late stage of pairing (>1 moth post-pairing), which would be unambiguously considered pair maintenance.

### Behavioral recordings

Recordings were made in a sound-attenuated test room, separate from the colony room. Partners were transported, one at a time, in a small covered cage to separate, individual cages each equipped with a tie-clip microphone and a piezo electric sensor attached to the perch (Figure 1). The transport of each partner took 2-3 min, thus partners were separated for 4-6 minutes during transportation. This brief separation period is sufficient to elicit a social reunion (greeting behavior). Partners were not always transported in the same order (some days the male was transported first and other days the female was transported first). Upon placement of the second partner in the cage, reunion behavior was recorded. All four channels (one movement and one acoustic channel for each partner) were recorded using a multi-channel Zoom recorder (F8). Thus, we were able to make single recordings with temporally aligned, individually identifiable acoustic and movement behavior from each partner.

We have previously demonstrated that the majority of activity during this behavioral assay occurs within the first few minutes (<5 min) (Prior et al., 2019). First, we minimized the stress of handling by habituating birds to the behavioral and transport procedure prior to the start of the experiment. Additionally, once in the testing room, birds were not handled and were allowed to enter the testing cage/exit the transport cage on their own, with the researcher present. Second, we habituated birds to the procedure prior to the start of experimentation to ensure bird that any changes in behavior would not be due to habituation to the paradigm. Habituation included 10 consecutive days of transport to the testing room (transport only = 4 days; testing room alone (individual recording) = 3 days; and social reunion with a same-sex flock mate = 3 days). The last day of habituation was the day before birds were moved to mixed-sex flocks. Third, we quantified behavior in our assay multiple times during each pairing stage (after the initial pairing). Despite that our behavioral assay clearly elicits socially-directed greeting/reunion behavior, we were concerned that such behavior during these brief behavioral assays would be easily influenced by other factors such as and individuals experience just prior to the assay (such as what was happening in the home cage prior to testing, and aspects of the transport). Finally, we included prior experience at the start of pairing as a factor in our analyses, as it may influence pairing dynamics.

### Scoring behavior during social reunions

#### Operational definition of calls and movements

Overall movement and calls rates were quantified for each partner during the first five minutes of the social reunion. Using an in-house written MATLAB program (written by E.S.) we automatically identified time-stamped movements for each partner (Prior et al., 2019). Following initial observations and pilot experiments we decided to group all movements together for three main reasons. First, individuals produced a small repertoire of behaviors. All of the behaviors we observed appeared to be related to the social interactions and included larger body movements (perch hops) and small movements (including head tilts, fluff ups, and wing movements). Note that our goal was to reliably elicit social interactions, so the cages were small and contained no additional stimulation or distraction. Second, it was difficult to disentangle discrete movements (both during observations and from the recording on the timewaveform) because large and small movements often occurred simultaneously or happened in rapid succession. Finally, we had no *a priori* reason to differentiate between large and small body movements.

Calls were semi-automatically classified to identify time-stamped calls for each partner (in-house written MATLAB program written by E.S.). An initial automatic classification identified all noises, including vocalizations and non-vocalizations. One researcher (N.H.P) manually classified all auditory events as non-vocalizations or calls from either Bird 1 or Bird 2 (left or right channel, respectively). Auditory events were manually classified based on visual assessment of the spectrogram and timewaveform and assigned to the appropriate channel based on the amplitude of the signal on the timewaveform. While call types were not distinguished in the current dataset, the large majority of calls produced were stack calls (with some distance calls). Stack calls are the most common call type used in this behavioral assay (Prior et al., 2019), are commonly used between mates outside of a breeding context (D’Amelio, 2018; Pietro Bruno D’Amelio et al., 2017b; Gill et al., 2015), and encode information on sex and individual identity (Pietro B. D’Amelio et al., 2017; Prior et al., 2018). Stack calls appear to be important for communicating information about movement of partners when they are separated by short distances (D’Amelio, 2018; Pietro Bruno D’Amelio et al., 2017b).

#### Operational definition of behavioral coordination

The coordination of movements and calling was quantified separately (Prior et al., 2019). First, the similarity in activity rates within a dyad were calculated (e.g. Similarity of Calling = Call Rate of Bird 1 – Call Rate of Bird 2 / Call Rate of Bird 1). Second, the temporal synchronization of social dynamics was estimated using sliding correlation coefficients (based on Pearson correlations) of the time-stamped list of movements and events that was generated for each recording. More specifically, the sliding correlation coefficients were calculated separately within the vocalizations and movements using MATLAB “corrcoef” function. The step size for the sliding correlations was chosen based on the natural temporal dynamics of the movements and calls which we assessed during preliminary observations and development of the programs. For calculation of calling synchrony, a 1 ms sliding correlation timestep was used. For calculation of movement synchrony, a 40 ms sliding correlation timestep was used. Inputs to the sliding correlation computations were two vectors of ones and zeros, with a one indicating presence of a movement or call during the sliding window (1 ms for calls and 40 ms for movements) and a zero indicating absence of a movement or call. The sensor signal power in each time step was computed. For statistical analyses we used the maximum Pearson’s correlation coefficient value (“corrcoef”) based on all possible temporal offsets.

#### Disentangling activity and coordination for calls and movements

The above approaches allow us to quantify two crucial components of the social reunion: activity and coordination. However, both mathematically and conceptually, the amount of activity and the coordination of activity may be related. Furthermore, the above approaches operationalize movement and calling separately but we would expect these two behaviors to be related. Therefore we used principal components analyses (PCAs) to determine whether we could disentangle activity from behavioral coordination, as well as to describe multi-modal behavioral profiles of these social interactions. We conducted two separate PCAs, one for females and one for males (function ‘prcomp’) because there was a significant effect of sex on activity (*see results)*. Four dependent variables (call rate, movement rate, sliding correlation of time-stamped calls, sliding correlation of time-stamped movements) were loaded into each PCA.

For females, the first two components explained 67% of behavioral variation (PC1 explained 36% and PC2 explained 31%). PC1 describes overall activity, with call rate and movement rate positively related (Loadings: Call Rate = −0.625; Movement Rate = −0.732, Sliding Correlation Coefficient of Calls = 0.185, Sliding Correlation Coefficient of Movements = 0.227), whereas PC2 describes coordination of activities with a positive relationship between call rate, the coordination of calling and the coordination of movements (Loadings: Call Rate = 0.444; Movement Rate = −0.018, Sliding Correlation Coefficient of Calls = 0.661, Sliding Correlation Coefficient of Movements = 0.605).

For males, the first two components explained 74% of the behavioral variation (PC1 explained 46% and PC2 explained 28%). PC1 describes overall activity and the coordination of calls with call rate and movement rate positively related (Loadings: Call Rate = −0.663; Movement Rate = −0.642, Sliding Correlation Coefficient of Calls = −0.372, Sliding Correlation Coefficient of Movements = 0.104), whereas PC2 describes coordination of activities with call rate and movement rate positively related (Loadings: Call Rate = 0.190; Movement Rate = 0.227, Sliding Correlation Coefficient of Calls = −0.502, Sliding Correlation Coefficient of Movements = −0.812).

### Statistical analysis

All statistical analyses were carried out in R (v.3.2.3, R Foundation for Statistical Computing). We used linear-mixed models (function lmer from the lme4 Package). For each model, prior to interpretation, we transformed data as necessary based on a visual assessment of the distribution of the residuals. All data presented in graphs is non-transformed.

The effect of pair bonding on activity levels (call rate and movement rate) of males and female partners during social reunions was assessed using linear-mixed models with Pairing Stage and Sex as fixed factors and Individual identity as a random factor (CallRate∼Sex*Pairing Stage+(1|BirdID)). The effect of pair bonding on the coordination of activities was assessed using linear-mixed models with Pairing Stage as a fixed factor and pair ID as a random factor (Sliding correlation coefficient calling∼ Pairing Stage+(1|PairID)). Similarly, the effect of pair bonding on multimodal principal components was assessed separately for males and females with Pairing Stage as a fixed factor and Individual ID as a random factor.

As we described above, we distributed the nine social reunion recordings based on key stages or pair bonding, rather than evenly throughout the month. For that reason, we chose to use Pairing Stage rather than Date as our primary variable. However, as a double check, the significant main effects that we report here for Pairing Stage are also present if we use Date as the primary factor. Additionally, because we had the expectation that prior experience at the time of pairing would influence social interactions, we subsequently ran linear-mixed models for key dependent variables including Prior experience as a fixed factor (Sliding correlation coefficient calling∼ Pairing* Prior Experience+(1|PairID).

## Results

### Activity levels of females and males during social reunions

While isolated (recorded individually in the testing room), both males and females were largely inactive (Figure 2). Eighteen out of the twenty individuals had a movement rate of <1/min (9 females and 9 males) and fifteen out of the twenty individuals had a call rate of <1/min (6 females and 9 males). We recorded each partner alone in the testing room as a control and it is not included in our statistical models: however, the low levels of activity during isolation emphasizes the extent to which we are able to elicit a social reunion. During the social reunions both call rate and movement rate were higher for females than males, regardless of the stage of pair bonding (Figure 2. Call Rate *χ*^2^ (1) = 4.65, P = 0.031; Movement Rate *χ*^2^ (1) = 5.72, P = 0.017).

**Figure 2:**
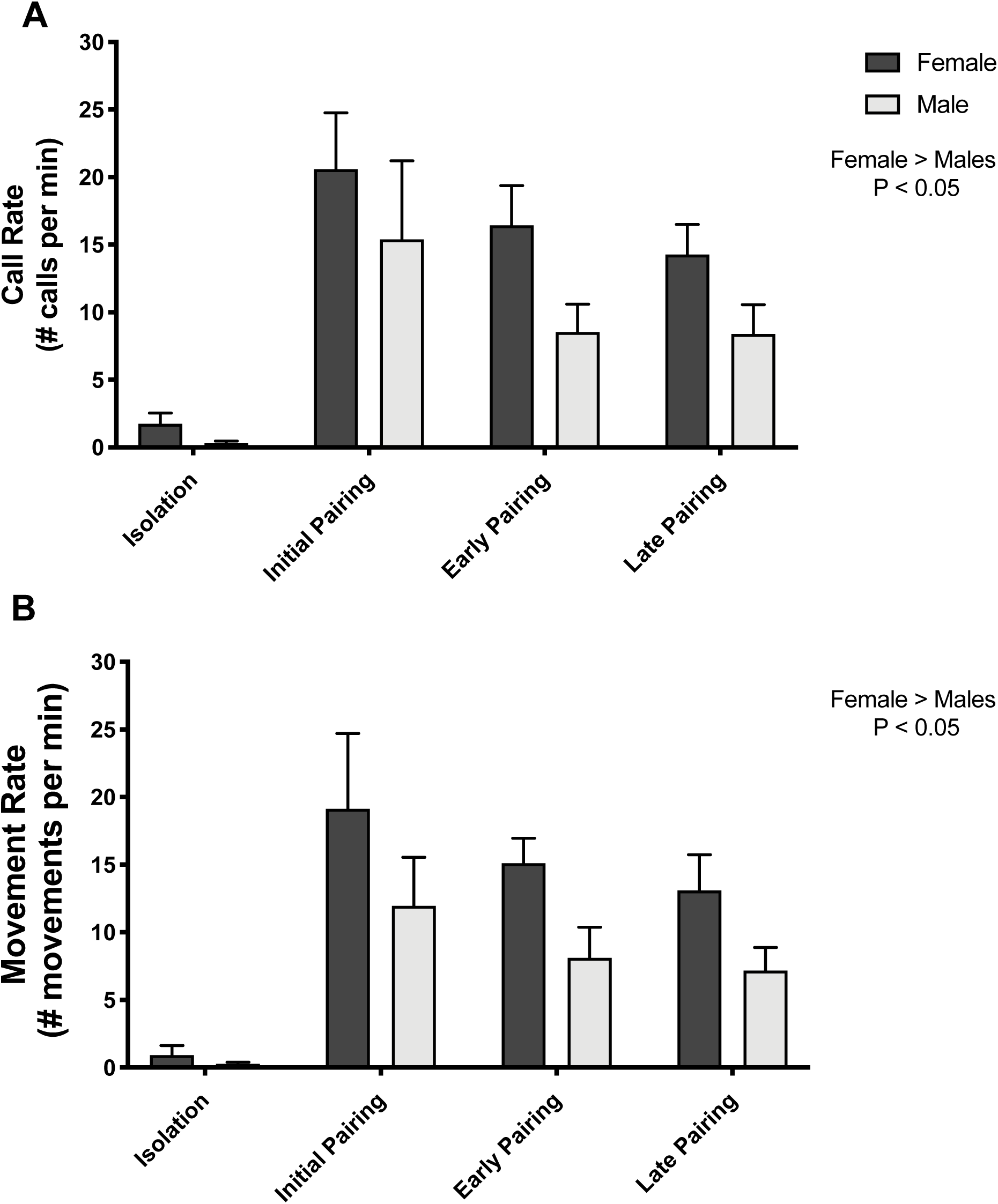
Overall activity rates, call rate (A) and movement rate (B), are shown for females and males tested in isolation and with their partner across the three pairing stages. Overall, females were more active than males. Additionally, male and female call rates were highest during the initial recording (Pairing Stage *χ*(2) = 8.35, P=0.015; Pairing Stage × Sex *χ*(2) = 0.677, P=0.713).

### Calling activity is modulated by pairing

For both males and females call rates were highest during the initial recording (Figure 2. Pairing Stage *χ*^2^ (2) = 8.35, P=0.015; Pairing Stage × Sex *χ*^2^ (2) = 0.677, P=0.713). This main effect was driven by a difference between the initial time point and the later timepoint (summary LMER t value = −1.971 P= 0.051. There is a similar pattern of decreased movement rate during later pairing, but this effect is not significant (Figure 2. Pairing Stage *χ*^2^ (2) = 4.52, P = 0.104; Pairing Stage × Sex *χ*^2^ (2) = 0.30, P = 0.861). The results from our PCA analyses further support the interpretation that the effect of pairing stage is specific to call, not movement, rate. For both males and females, PC1 represented a composite multimodal activity score (call rate and movement rate were positively correlated *see methods*). There was no effect of pairing stage on PC1 for females (*χ*^2^ (2) = 4.21, P=0.122) or males (*χ*^2^ (2) = 3.43, P=0.180).

### Pairing stage had no effect on the coordination of activities

There was no effect of pairing stage on coordination of activity for either calls or movements (Figure 3). Pairing stage had no effect on the percent difference in call rate (F:M) (*χ*^2^ (2) = 1.47, P = 0.481), nor on the sliding correlation coefficient of calling activity (*χ*^2^ (2) = 2.85, P = 0.240). Likewise, there was no main effect of pairing stage on the coordination of movements (percent difference *χ*^2^ (2) = 0.11, P = 0.944; sliding correlation coefficient *χ*^2^ (2) = 1.00, P=0.605). Again, the results of our PCA are consistent with the raw data. For both males and females PC2 represented a composite multimodal coordination score (the sliding correlation coefficient of calls and movements were positively correlated *see methods*). There was no effect of pairing stage on PC2 for females (*χ*^2^ (2) = 1.09, P=0.581) or males (*χ*^2^ (2) = 3.35, P=0.187).

**Figure 3:**
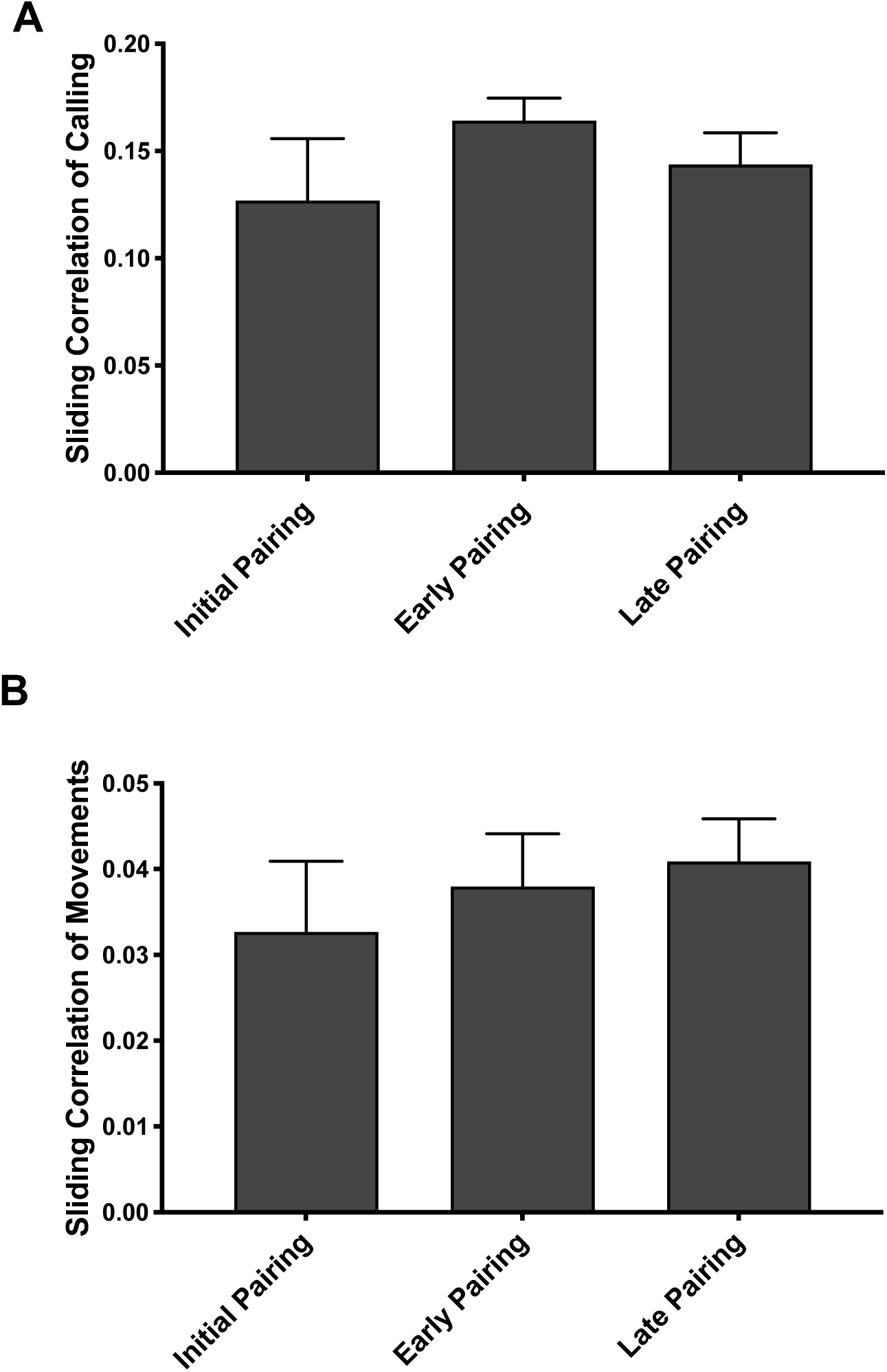
Pairing stage had no effect on the coordination of activities between males and females as quantified using the sliding correlation coefficient of time-stamped calls (A), and time-stamped movements (B), across the three pairing stages. As described in the text, there was also no effect of pairing stage on similarity of call rate and movement rate between partners.

### Additional factors that may influence social reunion

Pairing stage was the primary factor we investigated. However, we had the opportunity to also ask whether there was the potential for a relationship between two additional features of the partnership and behavior during the social reunion. First, we asked whether prior experience at the time of pairing may influence behavioral coordination. Indeed, there was a significant interaction between Experience and Pairing Stage on activity (Figure 4A-D. Calling Prior Experience *χ*^2^ (2) = 0.10, P=0.950; Pairing Stage × Prior Experience *χ*^2^ (4) = 17.44, P = 0.001; Movement: Prior Experience *χ*^2^ (2) = 0.20, P=0.903; Pairing Stage × Prior Experience *χ*2 (4) = 17.88, P = 0.001). There was also a significant interaction between Experience and Pairing Stage on calling synchrony (sliding correlation coefficient of calls: Prior Experience *χ*^2^ (2) = 3.31, P=0.191; Pairing Stage × Prior Experience *χ*^2^ (4) = 17.93, P = 0.001). However, there was no effect of prior experience on any other measure of the coordination of activities (Percent Difference in Calling: Prior Experience *χ*^2^ (2) = 2.03, P = 0.363; Pairing Stage × Prior Experience *χ*2 (4) = 7.01, P = 0.135; Percent Difference in Movement: Prior Experience *χ*^2^ (2) = 1.16, P = 0.561; Pairing Stage × Prior Experience *χ*^2^ (4) = 2.57, P = 0.633; Sliding Correlation of Movements: Prior Experience *χ*^2^ (2) = 0.30, P = 0.863; Pairing Stage × Prior Experience *χ*^2^ (4) = 4.49, P = 0.383).

**Figure 4:**
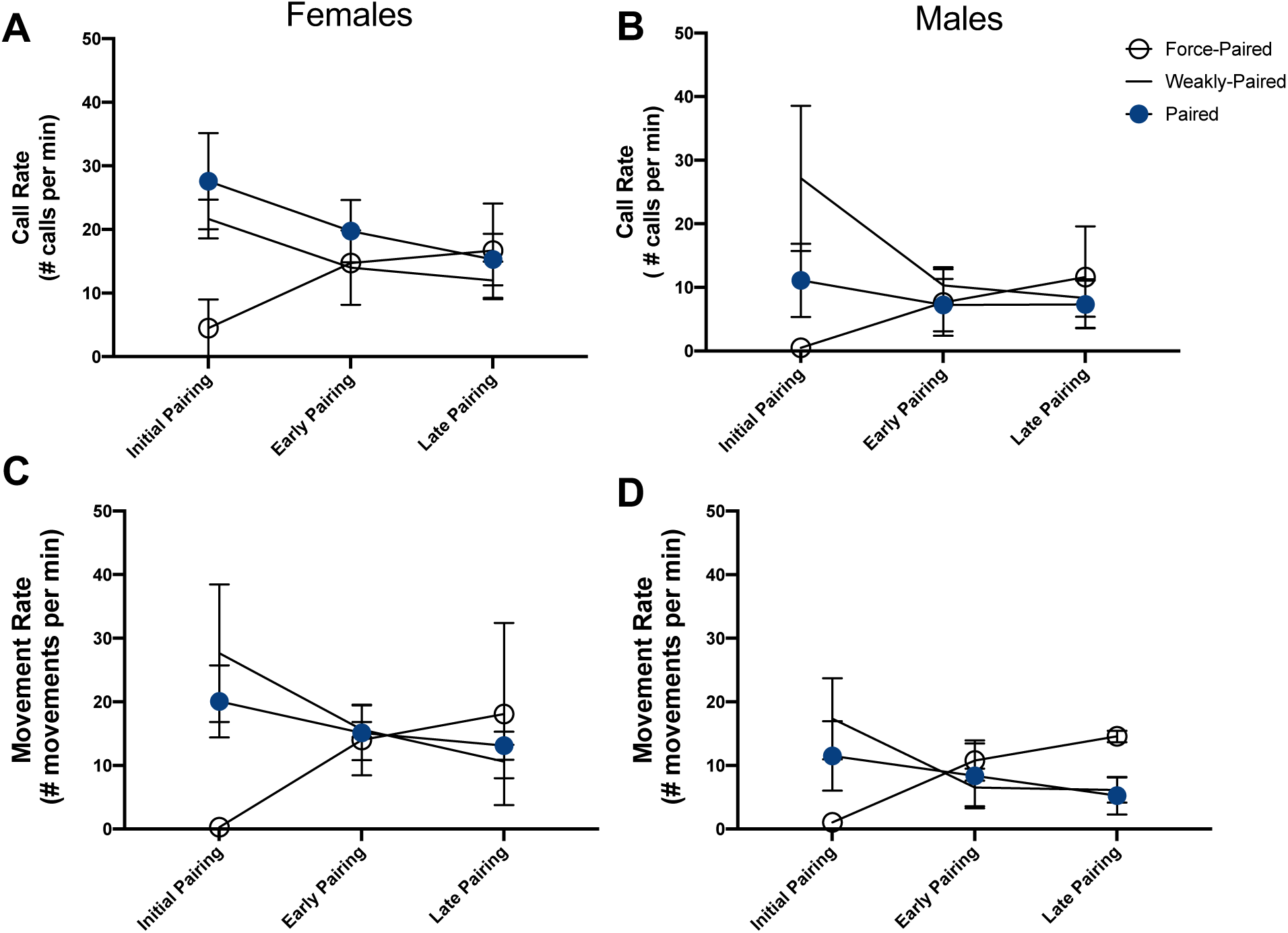
Effect of prior experience on activity (call rate A-B and movement rate C-D) for females (A and C) and males (B and D) across the three stages of pairing. Pairs differed based on the amount of time and prior experience they had when they were initially paired after the 72 hours being housed in mixed-sex flocks. Four pairs had clearly formed (paired). Another four pairs were created from individuals of the same mixed-sex flocks (weakly paired), with little evidence of pairing in the flocks. The final two pairs were composed of individuals that had no prior familiarity with each other (force paired). There was no effect of sex on either calling or movement, however there was a significant interaction between Prior Experience and Pairing Stage (Calling: *χ*(4) = 17.44, P = 0.001; Movement: *χ*(4) = 17.88, P = 0.001).

Second, we tested whether behavioral coordination during social interactions during early pairing influences breeding success. After the conclusion of this study, pairs had multiple opportunities to breed together. In the first breeding attempt (two months after the end of this experiment) all 10 pairs engaged in nest building behavior and laid eggs. Seven out of the 10 pairs went on to successfully fledge chicks (between 2-4 chicks per clutch). One of these only built a nest (a strongly paired dyad). The two other pairs laid eggs that did not hatch (a weakly paired and force paired dyad). There was no difference in calling synchrony or movement synchrony between pairs that successful fledged chicks (Sliding Correlation of Calls: *χ*^2^ (2) = 0.24, P = 0.621; Sliding Correlation of Movements: *χ*^2^ (2) <0.01, P = 0.978).

## Discussion

Here we describe the subtle changes that occur during brief social reunions between monogamous partners over the first month of pairing. The primary effect was elevated calling of both the male and female partner during the initial social reunion timepoint. Interestingly, we saw no change in coordination of calling or movement activity over the course of pairing. This is particularly notable because we have demonstrated previously that social reunions with novel conspecifics elicit less robust behavioral responses and that those interactions are less coordinated (Prior et al., 2019). Together this suggests that zebra finches are very rapidly able to develop patterns of interactions between familiar individuals.

Beyond the primary effect of pairing on calling activity, this study raises additional questions about what factors influence the social reunion of partners. First, across pairing stages there were differences between females and males in their respective contributions to the interaction. Overall, females were more active than males (having higher call rates and movement rates). Second, pairs differed in how long they had been paired and the nature of the courtship experience (prior experience). Our data suggests this prior experience influenced patterns of behavior (both activity and the coordination of activities) during the initial, but not subsequent reunions.

### Changes in pair-directed affiliative behavior over time

For species that maintain long-term social bonds, including humans, the ability to maintain such bonds is as important as the ability to form them. Importantly, there is evidence that the neurobiological mechanisms underlying the formation and maintenance of bonds are different (Aragona et al., 2006; Prior & Soma, 2015; Resendez et al., 2016; Smiley et al., 2012). There are many highly-marked affiliative behaviors that are associated with pair bonding, which can be useful in some species for identifying the point when pair bonds become established. Research investigating the changes in pairing over time has tended to parse these stages of pair bonding into discrete stages. In species such as zebra finches, where a mating event may not be needed for pair bond formation and individuals do not form a strong partner preference, it is particularly challenging to distinguish discrete changes.

Whereas the presence of highly-marked affiliative behaviors (e.g. clumping (close proximity), side-by-side perching facing the same direction, and allopreening (Black & Hulme, 1996; Reichard & Boesch, 2003; Zann, 1996)) are associated with the establishment of a bond, it is less clear how these behaviors change over time following initial pair bond formation. There is some evidence from both prairie voles and zebra finches that even after pair bond establishment, pairs continue to increase the amount of time they spend in close proximity (Pietro Bruno D’Amelio et al., 2017b; Scribner et al., 2020).

In zebra finches, D’Amelio and colleagues (2017) described the changes in social dynamics of new vs established zebra finch pairs in their home cage over a week. On the first day, newly paired zebra finches spent significantly less time in physical proximity (clumping) compared to the established pairs but groups were already more similar by the third day of pairing. This timeline is consistent with when we see most significant changes in social reunion behavior in the current study (within the first few days of pairing). However, the trend for a difference between new and established pairs in D’Amelio et al., 2017 was present throughout the week of recording. Similarly, in prairie voles time spent in close proximity between partners appears to increase over the first month of pairing (Scribner et al., 2020).

D’Amelio and colleagues (2017) also carefully described the effect of pair bonding on vocal interactions of new and established zebra finch pairs using continuous vocal recordings. On the first day of recording, newly paired individuals called less than predicted. Additionally, over the first week of pairing, calling dynamics between partners became more symmetrical: shifting from a scenario where one bird calls more and the other answers to a scenario where each partner calls and answers similarly. Importantly, even newly paired birds were clearly motivated to engage in call-response. It is striking that we see similar patterns emerging during the brief social reunion assay. The types of calling exchanges we elicited here are very similar to those elicited by D’Amelio et al., 2017, and are predominately made up of stack-stack calls and responses. This highlights the ethological relevance of our current social reunion test. Our current study combined with recent research highlights the need for a more comprehensive description of the subtle ways that social dynamics of partners change over time (Scribner et al., 2020).

### Monogamy and moment-to-moment behavioral coordination

Broadly, behavioral coordination across timescales is associated with gregariousness (Conradt & Roper, 2005; Focardi & Pecchioli, 2005) and is thought to increase social cohesion (King & Cowlishaw, 2009; Pays et al., 2007) and affiliative behavior (Sakai et al., 2010). Furthermore, behavioral coordination between two individuals has been shown to promote prosocial behavior (Ashton–James et al., 2007; Gueguen et al., 2009; Van Baaren et al., 2004) (reviewed in (Duranton & Gaunet, 2016)), which we would expect to reflect the presence of affiliative bonds. Even on extremely acute timescale, such as what we are focused on here, there is evidence in humans that social or interactional synchrony promotes the formation and reinforcement of affiliative bonds (Feldman, 2007; Feldman & Eidelman, 2007; Feldman et al., 2011). In songbirds specifically, there has been extensive research investigating the function of behavioral coordination in monogamous pairs particularly as it relates to breeding success and the coordination of biparental care (I. C. A. Boucaud et al., 2016; Mariette & Griffith, 2012, 2015; van Rooij & Griffith, 2013). Interestingly, brief social interactions at the nest, only a few minutes long appear to be essential for the coordination of parental duties across many species (review Prior 2020). In zebra finches specifically, there are several lines of evidence demonstrating that parental duties are actively coordinated during interactive calling exchanges (I. Boucaud et al., 2016; Boucaud et al., 2017; Elie et al., 2010; Villain et al., 2016). During incubation, female calling behavior predicts whether or not the male will relieve her and begin incubation (Boucaud et al., 2017). Experimentally preventing the male from returning to the nest to relieve the female, causes her to modify calling behavior which subsequently predicted the timing of the females time off the nest (I. C. A. Boucaud et al., 2016). Experimental evidence from other avian species suggests that the coordination of parental behavior is an emergent consequence of the behavioral interactions between partners, not simply a summation of both partners contributions (Ball & Silver, 1983). How partners develop such patterns of communication and what makes partners good communicators remains an open question.

It is possible that the patterns of a partner’s communication during mundane social interactions lays the foundation for these marked-moments. After the end of our current experiment, all the pairs were given the opportunity to breed. Seven out of 10 pairs eventually went on to successfully fledge chicks, although there were no differences in behavioral coordination of the pairs that successfully fledged chicks and those that did not. The fact that we saw no relationship between behavioral coordination and breeding success is not surprising given our very low sample size (3 vs 7) as well as the amount of time after the last social reunion recording and breeding. Furthermore, we would expect that there are nuances beyond temporal synchrony alone which are important for the development of these social interactions. Future research will directly test the relationship between moment-to-moment coordination of activities and the coordination of parental behavior in order to further elucidate the function of behavioral variation in these brief interactions.

### Effect of familiarity and prior experience on social reunions

An intriguing potential confound exists when examining the long-term effects of pair bonds on social dynamics: to what extent can the nature of the social relationship (formation of a monogamous pair bond) be disentangled from familiarity and shared social experience. Using the same social reunion behavioral assay, we recently demonstrated that familiarity itself, not pair bonding per se, influences social reunion behavior (Prior et al., 2019). When we compared reunion behavior between different social dyads (monogamous partners, familiar same-sex dyads, familiar opposite-sex dyads, novel same-sex dyads, and novel opposite-sex dyads), we found that both activity levels and the coordination of activity was higher in familiar social dyads (Prior et al., 2019). In our current study, only the two force paired dyads (paired for ∼4 hours), appeared similar to the novel dyads described previously. Thus, we can assume that pairs were able to familiarize and stabilize their degree of coordination very quickly, between 4 – 72 hours. Combined, we interpret the results of these two studies as demonstrating an effect of prior social experience on behavioral coordination and patterns of behavior during brief social interactions.

Over the course of pair bonding, the fact that coordination is maintained but the activity decreases could be evidence that the strength of established bonds is positively correlated with the length of social exchanges required during. Perhaps well-established partners require ever-briefer, and less intense, social exchanges. Perhaps it is less the extent of coordination, but rather the extent of the effort it takes to coordinate that reflects pair bond strength (although prior experience had no lasting effect on behavioral profiles beyond behavior during the initial courtship timepoint). Despite behavioral coordination/ interactional synchrony being influenced by social bonding; the processes by which this happens may be a shared biological foundation of social alignment, rather than a pair bonding process specifically.

### Conclusion

Here we show one way that mundane social interactions is affected by pair bonding, likely via shared prior social experience. Broadly speaking, this is consistent with the idea that social relationships are a culmination of repeated social interactions between familiar individuals. Understanding how brief social exchanges are modulated by experience and social bonding may provide an entry point to describe the wide diversity in the phenotypes of social relationships.

## Acknowledgements

We would like to thank the entire Ball/Dooling lab for help with animal husbandry, data collection, and discussion of results. An earlier version of these results was presented at the Animal Behavior Society meeting in Chicago, IL in July 2019. We are grateful for all of the feedback we received there. For feedback on this manuscript, we thank Dr. Matthew D. Taves and Dr. Benjamin A. Sandkam.

## Ethics Statement

This work was conducted in accordance with Association for the Study of Animal Behaviour (ASAB) guidelines and was approved by the Institutional Animal Care and Use Committee (IACUC) (R-15-09), University of Maryland, College Park.

